# Universal signatures of transposable element compartmentalization across eukaryotic genomes

**DOI:** 10.1101/2023.10.17.562820

**Authors:** Landen Gozashti, Daniel L. Hartl, Russell Corbett-Detig

**Affiliations:** Department of Organismic and Evolutionary Biology, Harvard University, Cambridge, MA, USA; Museum of Comparative Zoology, Harvard University, Cambridge, MA, USA; Department of Biomolecular Engineering, University of California Santa Cruz, Santa Cruz, CA, USA; UC Santa Cruz Genomics Institute, University of California Santa Cruz, Santa Cruz, CA, USA

## Abstract

The evolutionary mechanisms that drive the emergence of genome architecture remain poorly understood but can now be assessed with unprecedented power due to the massive accumulation of genome assemblies spanning phylogenetic diversity ^1,2^. Transposable elements (TEs) are a rich source of large-effect mutations since they directly and indirectly drive genomic structural variation and changes in gene expression ^3^. Here, we demonstrate universal patterns of TE compartmentalization across eukaryotic genomes spanning ∼1.7 billion years of evolution, in which TEs colocalize with gene families under strong predicted selective pressure for dynamic evolution and involved in specific functions. For non-pathogenic species these genes represent families involved in defense, sensory perception and environmental interaction, whereas for pathogenic species, TE-compartmentalized genes are highly enriched for pathogenic functions. Many TE-compartmentalized gene families display signatures of positive selection at the molecular level. Furthermore, TE-compartmentalized genes exhibit an excess of high-frequency alleles for polymorphic TE insertions in fruit fly populations. We postulate that these patterns reflect selection for adaptive TE insertions as well as TE-associated structural variants. This process may drive the emergence of a shared TE-compartmentalized genome architecture across diverse eukaryotic lineages.

## Main

The evolutionary forces that drive the emergence of eukaryotic genome architecture remain largely obscure. However, the breadth of annotated genome assemblies spanning a large portion of eukaryotic diversity now makes it possible to identify factors that shape genome evolution ^1,2^. Transposable elements (TEs) are ubiquitous parasitic genetic elements capable of dispersing copies across host genomes and function as major drivers of molecular and phenotypic variation^3^. Despite their selfish behavior, TEs often contribute to adaptation at individual loci as well as large-scale reshaping of gene regulatory networks resulting in adaptive novelty ^3–5^. Furthermore, TEs passively shape the distribution of genomic structural variants (including gene duplications and deletions) due to their tendency to facilitate nonallelic homologous recombination (NAHR) as well as their susceptibility to double stranded breaks during replication, with TE-rich genomic regions displaying a greater propensity for structural variation than TE-poor regions ^6–8^ (Figure 1A). TEs are non-randomly distributed across many genomes, and often show massive enrichment in specific compartments ^9–13^. Although TE-rich compartments are usually depleted in genes likely due to the deleterious effects of TE-associated variation, some genes show enrichment for these regions ^9,10,13,14^. This “TE compartmentalization” of specific genes may arise as a consequence of direct selection on TEs that generate adaptive variation or passively when TEs create the preconditions necessary for adaptive structural rearrangements ^5,10,13,15–18^. Here, we demonstrate consistent patterns of TE compartmentalization across 1.7 billion years of eukaryotic evolution. This implies that natural selection drives emergent broad-scale properties of genome architecture.

**Figure 1:**
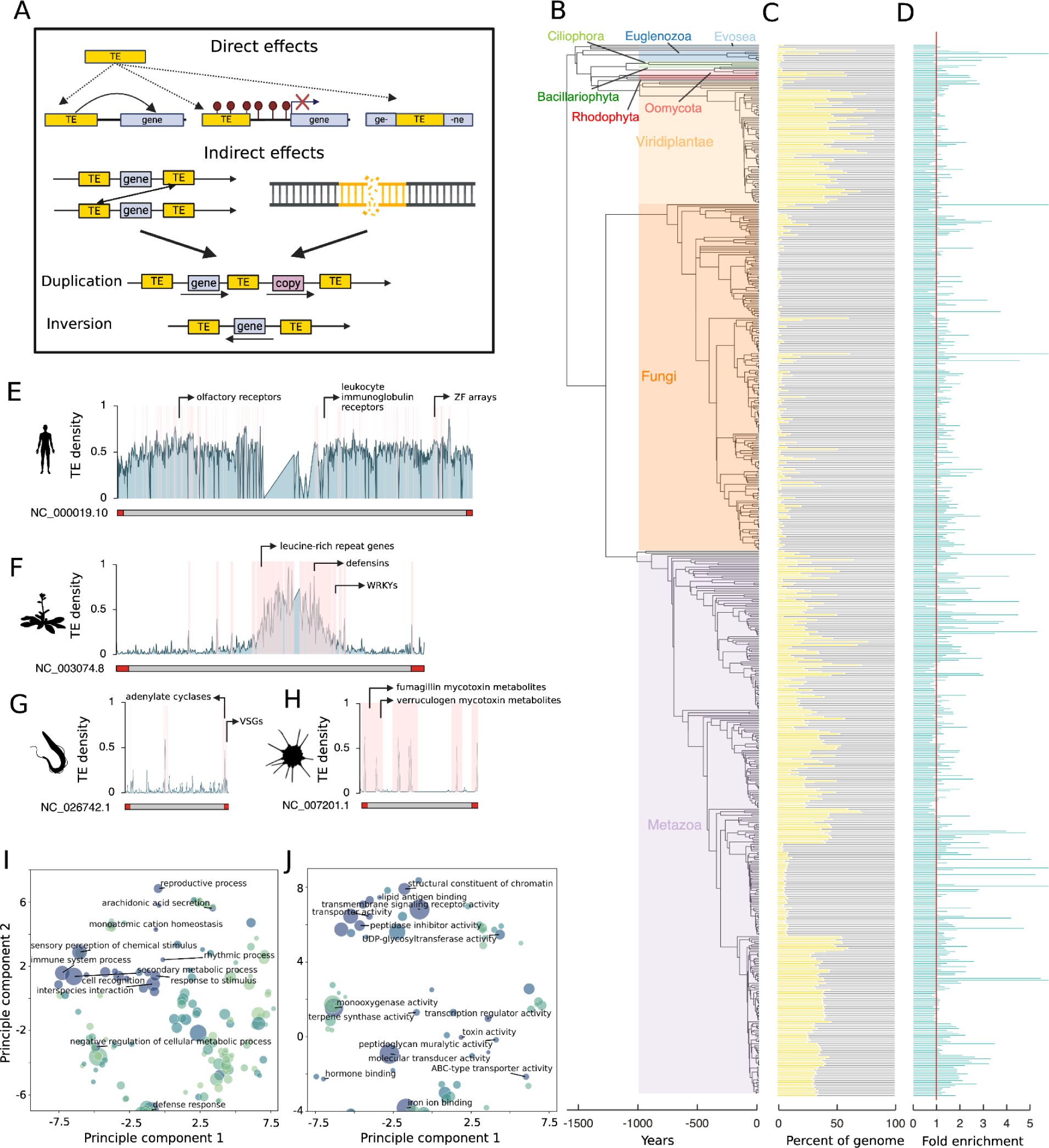
(**A**) Mutational effects of TEs. TEs can facilitate ectopic recombination and double stranded breaks resulting in genomic structural rearrangements such as duplications or inversions. TEs can also affect gene expression by distributing regulatory elements or altering heterochromatin environments. **(B)** Phylogeny of surveyed eukaryotic species. Branch lengths in millions of years were obtained from Timetree ^21^. (**C**) Barcharts show proportions of the genome occupied by TEs for each species. **(D)** Fold enrichment of TE-compartmentalized genes for multigene families. Values above the red line (placed at one) enriched. **(E-H)** Ideograms displaying TE density for (**E**) *Homo sapiens* chromosome 9 (Refseq NC_000019.10), (**F**)

### TEs colocalize with multigene families involved in specific functions

We developed a pipeline to search for patterns of colocalization between transposable elements and gene families involved in specific functions across diverse eukaryotes. To do this, we first generated *de novo* TE libraries and annotations as well as genome-specific gene ontology (GO) databases for 1,068 annotated genomes available through Refseq or Genbank (Supplementary Table 1; see methods). We performed extensive systematic and manual filtering for genome and GO database quality, resulting in a final set of 732 genomes (Supplementary Table 2). Then, for each species we extracted the 50 kb flanking regions of each gene and identified genes whose flanks displayed the top 90th percentile of TE content, which we refer to as “TE-compartmentalized genes” (see methods). We searched for commonalities among TE-compartmentalized genes using GO enrichment tests (Supplementary Data 1; Supplementary Tables 3-6). Due to massive variation in GO annotation extent and quality across considered species, with genes in some species showing annotations as specific as “evasion of host immune system” and others showing more general annotations such as “protein binding” or “membrane”, we also conducted extensive literature reviews to illuminate patterns obscured by less specific annotations (Supplementary Data 1).

Our results reveal consistent patterns of enrichment for multigene families involved in specific functions across 1.7 billion years of eukaryotic evolution notwithstanding massive variation in TE content (Figure 1B-C). In nonpathogenic species, TE-compartmentalized genes include those involved in sensory perception, environmental interaction and defense, whereas for pathogenic species, genes include those involved in pathogenicity. Furthermore, TE-compartmentalized genes are enriched for multigene families, with ∼83% of considered species showing as much as a 20 fold increase in multi-gene family representation (genes with >9 homologs) within TE-compartmentalized genes compared to other genes (Binomial test, P<0.000001; Figure 1D; Supplementary Tables 7-8). Remarkably, these patterns persist despite the lack of broad-scale synteny ^19^, multiple independent origins of gene families and functions ^20^, and vastly different repeat landscapes across divergent eukaryotes ^3^ (Figure 1C, E-H).

We hypothesize that these and other shared patterns reflect a combination of direct and indirect selection on TE-associated variation affecting genes under strong positive selective pressures. Specifically, TE insertions alter regulatory landscapes of neighboring genes (*e.g.,* by introducing enhancer binding sites or triggering host-mediated silencing). Additionally, TEs facilitate genomic structural rearrangements resulting in new gene duplications and deletions.

*Arabidopsis thaliana* chromosome 3 (Refseq NC_003074.8), (**G**) chromosome 9 of the trypanosome parasite, *Trypanasoma brucei* (Refseq NC_0267642.1), and **(H)** chromosome 8 of the pathogenic fungi, *Aspergillus fumigatus* (Refseq NC_007201.1) across 50 kb windows. TE-compartmentalized gene positions are highlighted in pink and the respective genomic positions of selected TE-compartmentalized gene families are labeled for each species. Red segments at the ends of chromosomes denote subtelomeric regions. (**I-J**) Multidimensional scaling results for enriched (**I**) Biological Process and (**J**) Molecular Function gene ontology (GO) terms for TE-compartmentalized genes across all surveyed species (Supplementary Tables 9-10). Each point represents a GO term, and point size and color correspond to enrichment frequency across species. Terms with dispensability metrics less than 0.0008 and 0.04 are labeled for Biological Processes and Molecular Functions respectively to prevent overcrowding.

### Enrichment for genes involved in sensory perception, environmental interaction, defense and pathogenicity

Our survey demonstrates remarkable patterns of enrichment for TE-compartmentalized genes involved in sensory perception, environmental interaction, pathogenicity and defense across the eukaryotic tree of life. This signal remains even when we aggregate our results across this exceptional diversity. Dimensionality reduction on aggregated enriched GO terms for “Biological Processes” reveals the strongest patterns for terms such as “immune system process”, “response to stimulus”, “interspecies interaction” and “secondary metabolism” (Figure 1I; Supplementary Table 9). Similarly, abundantly enriched “Molecular Functions” include “transmembrane signaling receptor activity”, “toxin activity” and “antigen binding” (Figure 1J; Supplementary Table 10). “Cellular Component” results generally reflect these patterns, showing enrichment for terms broadly associated with environmental interaction and excreted metabolites, such as “extracellular space” (Extended Data Figure 1; Supplementary Table 11).

### Sensory perception and environmental interaction

TE-compartmentalized genes show enrichment for sensory perception and environmental interaction. In metazoans, G protein-coupled receptors are enriched in species spanning from mouse (*Mus musculus*) to placozoans (*Trichoplax adhaerens*). GCPRs such as olfactory receptors represent some of the largest gene families in metazoans, play important roles in sensory perception and evolve rapidly in copy number due to positive selection (Figure 1E; Supplementary Tables 5-6) ^22,23^. Olfactory receptors more specifically are enriched in 79% of species with olfactory receptor annotations. Primarily sessile organisms such as land plants and fungi, and single cell organisms such as red algae secrete secondary metabolites to interact with their environments ^24,25^. Like GCPRs, genes involved in secondary metabolism and extracellular interactions are large gene families that evolve adaptively ^26–29^. In *Arabidopsis thaliana* for example, TE-compartmentalized genes include WRKY transcription factors, which play important roles in response to abiotic and biotic stresses and show patterns of rapid expansion and diversification across plant species (Figure 1F) ^30^. More broadly, genes involved in secondary metabolic processes and/or extracellular interactions are enriched in ∼64-75% of surveyed land plants and fungi (Figures 1I-K; Supplementary Tables 5-6).

### Defense

TE-compartmentalized genes also show enrichment for functions involved in defense in diverse eukaryotes. In the human genome, we observe enrichment for MHC complex genes as well as arrays of killer cell and leukocyte immunoglobulin receptors (Figure 1E). These innate and adaptive immunity genes are also among the most rapidly evolving genes in metazoans due to strong selective pressures from competing pathogens^31,32^. More broadly, 66% of vertebrates show enrichment for “MHC complex” genes (Supplementary Data 1; Supplementary Tables 5-6). We observe similar patterns for zinc finger (ZF) genes in metazoans, despite their poor annotation in most genomes (Figure 1E). ZFs are the most abundant transcription factors in metazoans and evolve rapidly under strong selective pressure due important for host defense against transposable elements (Figure 1E) ^33^. In plants such as *Arabadopsis thaliana*, we also find vital gene families for plant defense such as RING protein genes, leucine-rich repeat genes (NLRs), and defensins, consistent with previous reports (Figure 1F) ^10,18^. Consistent with observed trends for chordates, over 78% of land plants show enrichment for “defense response” in addition to other more specific terms related to immunity such as “terpene synthase activity” (Supplementary Data 1; Supplementary Tables 5-6).

### Pathogenicity

Pathogens frequently exhibit rapidly evolving “accessory genomes,” which harbor important gene families for pathogenesis and host adaptation. In the “two-speed” genomes of some fungal and oomycete plant pathogens, TEs colocalize with these gene families and play important roles in driving their dynamic evolution under positive selection ^10,34,35^. Our pipeline recovers expected patterns for species with known “two-speed” genome architectures, such as the potato blight-causing *Phytophthora infestans*, in which TE-compartmentalized genes are enriched for terms such as “modulation by symbiont of host programmed cell death” (Q<0.0001; Supplementary Table 4) ^36^. However, our results reveal similar trends in additional diverse pathogens. These include the trypanosome human parasite, *Trypanasoma brucei*, and the fungal human pathogen, *Aspergillus fumigatus*. TE-compartmentalized genes in *T. brucei* show up to nine-fold enrichment for functional terms like “evasion of host immune response” (Q<0.0001) and include variant surface glycoproteins and adenylate cyclases crucial for response to host defenses (Figure 1G; Supplementary Table 4) ^37,38^. In *A. fumigatus*, TE-compartmentalized include all annotated genes involved in verruculogen and fumagillin metabolism, mycotoxins linked to pathogenicity (Figure 1H; Supplementary Tables 3-4) ^39,40^. Gene ontology terms clearly related to host interactions and pathogenicity are poorly annotated for many species.

Nonetheless, 81% of species with “response to host immune response” gene ontology annotations show enrichment for TE-compartmentalized genes. Furthermore, many pathogens show enrichment for extracellularly secreted gene products as well as various enzymes and biosynthesis processes, as revealed by manual inspection ^10,11,41–44^.

### TE-compartmentalized gene families exhibit molecular signatures of positive selection

We hypothesized that TE-compartmentalized gene families should show evidence for strong positive selection at the nucleotide level, as reported for a subset of lineages where dynamic gene family evolution is associated with TEs^10,12,27,44–46^. To evaluate this prediction across eukaryotes, we tested for molecular signatures of positive selection for all gene families with at least 10 genes in each considered species (Supplementary Table 12; see methods). TE-compartmentalized gene families show an increased proportion of sites with signatures of positive selection compared to other genes (P(positive selection) > 0.95; Figure 2A, two-tailed binomial test, P< 0.001; Supplementary Table 12). Importantly, TE-compartmentalized gene families also show increased proportions of sites with signatures of constraint (P(constraint) > 0.95; Figure 2B; two-tailed binomial test, P=0.007; Supplementary Table 12), suggesting that observed patterns of increased TE content for TE-compartmentalized genes cannot be explained entirely by relaxed selection ^31,47^. Notably, we observe these trends regardless of taxonomic binning, even when highly divergent eukaryotic lineages are analyzed independently. These results suggest that TE-compartmentalized gene families evolve under positive selection.

**Figure 2:**
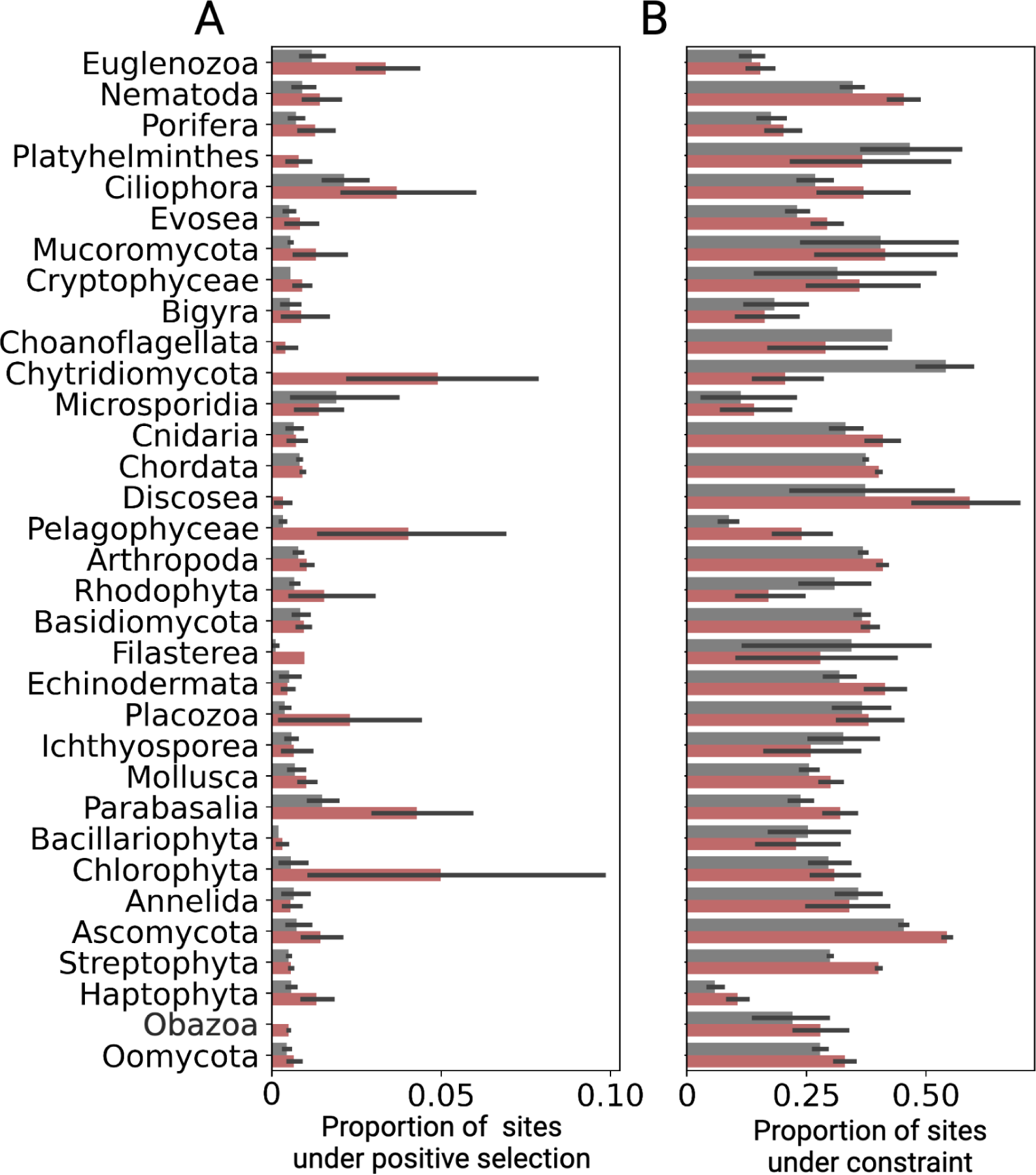
**(A-B)** Proportion of sites displaying significant signatures of **(A)** positive selection (P(positive selection) > 0.95) and **(B)** selective constraint (P(constraint) > 0.95) for TE-compartmentalized gene families (red) and other gene families (gray) in diverse eukaryotic taxonomic groups. Signatures of selection were predicted using FUBAR ^48^.

### TE-compartmentalized genes show diverse chromosomal distributions

Signatures of genomic compartmentalization are not constrained to specific regions along chromosomes. Genomic distributions of TEs vary tremendously across species, with some chromosomes showing increased TE content in subtelomeric regions and others showing contrasting patterns in which TEs cluster around centromeres ^9,34,49^. In many species, TE-compartmentalized genes show a bias towards subtelomeric regions, which are known to undergo frequent structural variation and serve as a cradle of adaptation in several lineages (Extended Data Figure 2; two-tailed permutation test, alpha = 0.05, N = 1000) ^34,49^. However, in other species we observe the opposite pattern, in which TE-compartmentalized genes are depleted from subtelomeric regions, suggesting that TE compartmentalization can arise irrespective of chromosomal position (Extended Data Figure 2). Importantly, we also find little evidence for consistent trends of association between TE-compartmentalized genes and recombination rates, or TE-compartmentalized genes and GC content, suggesting that these genomic features alone cannot explain observed patterns (two-tailed binomial test, P=0.151 and P=0.296 respectively; Extended Data Figures 3-4; see Methods).

### Positive selection might contribute to TE compartmentalization in populations

Positive selection might contribute to the origin and maintenance of TE compartmentalization within species. We used publicly available genomic data to analyze patterns of polymorphism in fruit fly (*D. melanogaster*) populations. Specifically, we compared the allele frequency spectrum of TE variants and single nucleotide variants (SNVs) for TE-compartmentalized genes and other genes across 50 long-read based fruit fly genomes (Supplementary Table 13). Interestingly, TE-compartmentalized genes exhibit an excess of high-frequency alleles for TE variants in their flanking regions, but not for SNVs in their coding regions (χ^2^ test P<0.001, Figure 3A). In fact, we observe a depletion of high-frequency SNVs in TE-compartmentalized genes compared to other genes (χ^2^ test P<0.001, Figure 3B-C). Assuming that most TE insertions around genes are deleterious or neutral, this pattern suggests that selection favors TE variants around TE-compartmentalized genes. Thus, positive selection might contribute to the evolution of TE compartmentalization.

**Figure 3:**
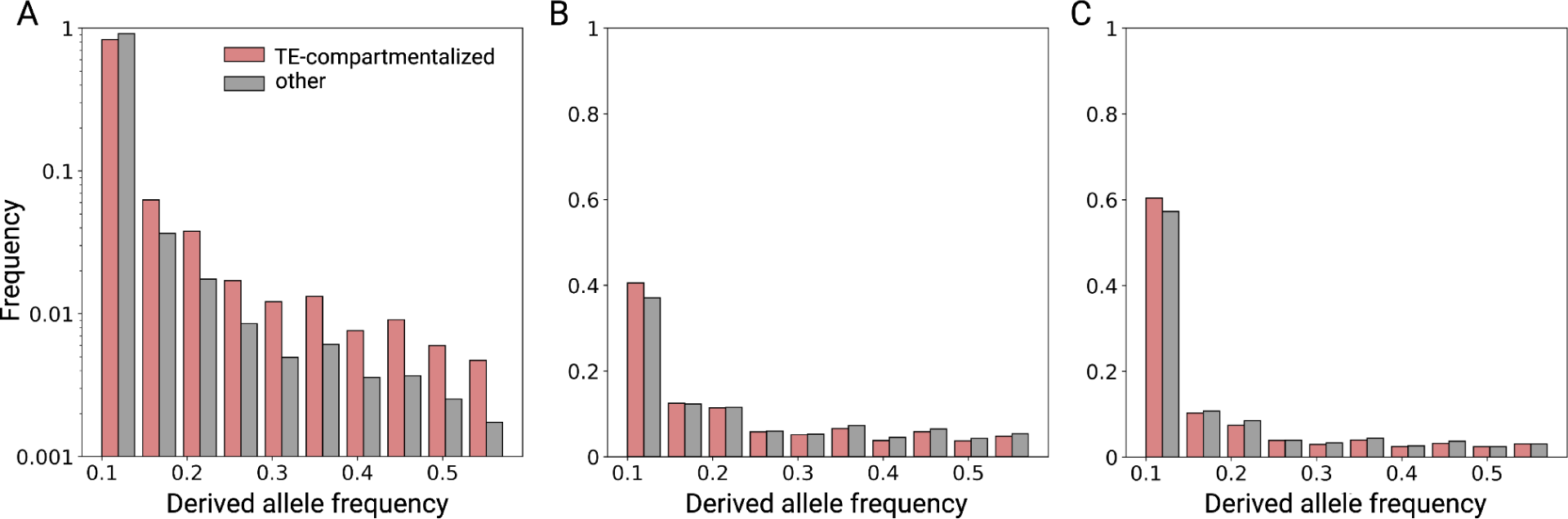
Folded allele frequency spectra for **(A)** TE variants in gene flanking regions, as well as **(B)** synonymous and **(C)** nonsynonymous SNVs in coding regions for TE-compartmentalized genes (red) and other genes (gray).

### TEs with the largest predicted mutagenic effects drive compartmentalization signatures in many species

TEs with the largest predicted molecular impacts are the most enriched in repeat hotspots surrounding rapidly evolving gene families. Longer TEs such as long interspersed nuclear elements (LINEs) and long terminal repeat (LTR) retrotransposons are more likely to generate genomic rearrangements and influence gene expression than shorter TEs such as short interspersed nuclear elements (SINEs) ^3,50^. We find that LINE-compartmentalized genes and LTR-compartmentalized genes show significantly different functional-enrichment profiles from SINE-compartmentalized genes for ∼79% and ∼75% of enriched functional terms respectively (Supplementary Tables 14-16; see Methods). Furthermore, LINEs and LTRs are the main contributors to compartmentalization signatures in species where LINEs, LTRs and SINEs are all present. For example, LINE-compartmentalized genes are enriched for olfactory receptors in ∼90% of chordates, whereas olfactory receptors are mostly underrepresented in SINE-compartmentalized genes (Supplementary Data 2). Similarly, LINE and LTR-compartmentalized genes are enriched for secondary metabolism in >74% of TE-containing ascomycete fungi, while SINE-compartmentalized genes are underrepresented for secondary metabolism in most ascomycete species (Supplementary Data 2). Although differences in TE insertional preference could also contribute to these patterns, the enrichment of longer TEs around rapidly evolving gene families is consistent with a model in which selection favors large-affect mutations driving dynamic gene family evolution.

### The TE compartmentalized plasticity model for genome evolution

The origins of eukaryotic genome architecture remain a fundamental question. Theory ^15^ and empirical analysis ^51,52^ show that genomes can restructure themselves due to the beneficial effects of genomic structural plasticity, resulting in the colocalization between TEs and gene families for which copy number variation is beneficial. Fungal plant pathogens similarly illuminate extreme examples of TE compartmentalization wherein genomes show bimodal distributions of TE content and TE-rich compartments house genes involved in pathogenicity^10–12^. Here, we demonstrate remarkable patterns of convergence across eukaryotic lineages spanning 1.7 billion years of evolution, in which TE-compartmentalized genes primarily represent multigene families involved in the same functions and evolve under positive selection. While relaxed purifying selection is a plausible alternative that may account for some of the excess of TEs observed for TE-compartmentalized genes ^18^, the lack of evidence for reduced constraint on these genes at the molecular level in diverse lineages as well as the excess of high-frequency derived TE insertions around such genes in fruit fly populations, is consistent with a role for positive selection.

We propose a compartmentalized plasticity model for genome evolution, wherein selection favors the colocalization between TEs and gene families for which rapid evolution is advantageous (Figure 4). Specifically, while TEs likely to impact gene expression or genome stability are generally depleted around essential genes due to their deleterious effects ^3,18,53,54^, selection may occasionally favor TE-associated mutations affecting rapidly evolving gene families. Over time, recurrent selective sweeps on TEs and TE-associated structural variants affecting these genes could result in TE compartmentalization. Subsequent TE compartmentalization functions as a positive feedback loop, promoting higher rates of variation and potential adaptive novelty for these gene families. These results illuminate a major universal emergent property of eukaryotic genome evolution.

**Figure 4:**
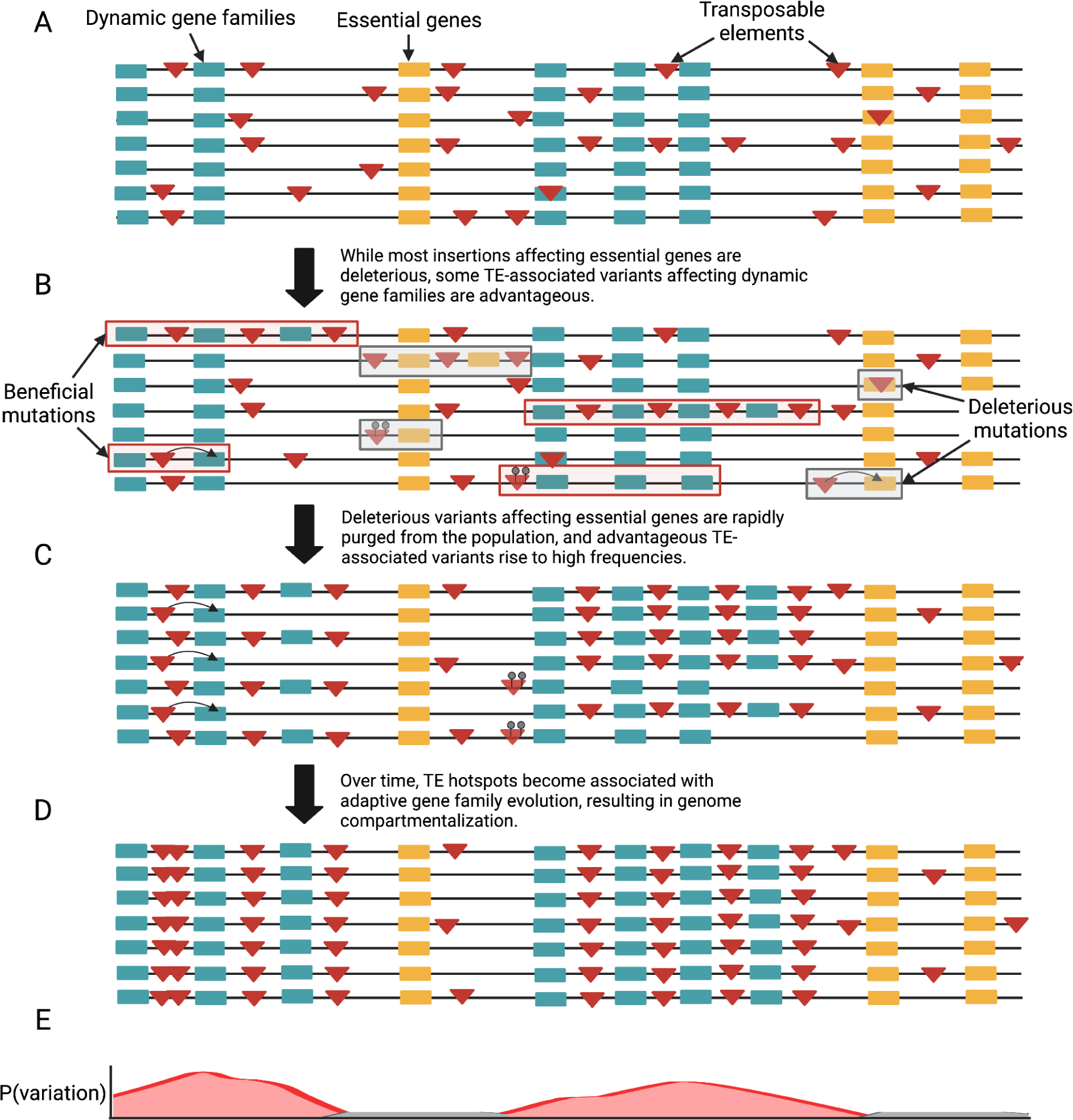
Compartmentalized plasticity model for genome evolution. Each line represents a haplotype within a population. Green boxes and yellow boxes delineate labile genes and essential genes respectively. Transposable elements are shown as red triangles. (**A**) Transposable elements insert semi-randomly across the genome. (**B**) Most TE insertions affecting essential genes (gray boxes) are deleterious. Some TE-associated variants affecting dynamic genes (red boxes) are advantageous. (**C)** Deleterious variants are rapidly purged from the population, whereas advantageous variants rise to high frequencies. (**D**) Over time labile genes colocalize with TEs, forming TE-rich compartments. (**E)** This colocalization functions as a positive feedback loop, since TE-compartmentalized genes exhibit an increased probability for future TE insertions and associated structural and regulatory variation.

## Methods

### Retrieving relevant genomic data

We downloaded all genome assemblies and annotations from NCBI ^55^ (Accessed February 20, 2020). We only considered genomes with RefSeq ^56^ assemblies for quality control purposes.

However, we opted to use GenBank ^55^ assemblies and annotations when GenBank annotations had a greater number of genes with ontology annotations. We downloaded functional annotations for each species from Uniprot ^57^. Accessions and metadata for all genomes used for the TE compartmentalization survey are displayed in Supplementary Table 1. We also downloaded additional *D. melanogaster* genomes from genbank for population analyses. Genome accessions for these samples are listed in Supplementary Table 13.

### Phylogenetic Display Items

The phylogeny in Figure 1 was retrieved from TimeTree <u>(Kumar et al. 2017)</u>.

### Repeat mining and annotation

To analyze patterns of repeat distributions across species, we first annotated transposable elements in each considered genome. We employed RepeatModeler (version 2.0.2) ^58^ to generate species-specific TE libraries *de novo* for each genome. Then, we processed each library to remove high copy number genes not associated with transposable elements. To do this, we used BLAST ^59^ to identify homologous sequences between candidate TEs and each genome’s transcript library as well as TE-related genes available from RepeatMasker. Then, we removed candidate TEs displaying strong homology (e-value < 1e-25) to species transcripts, ignoring those which showed homology to known TE genes. We used RepeatMasker (version 4.1.2) ^58^ to annotate TEs and simple repeats in each genome using each respective *de novo* repeat library as input, with the flags -*s -no_is -u -noisy -html -xm -a -xsmall* and resolved overlapping annotations using RM2Bed.py (https://github.com/rmhubley/RepeatMasker/blob/master/util/RM2Bed.py) with the *-o higher_score* flag. All TE libraries are being deposited to the DFAM database ^60^ following the first submission and will be finalized prior to formal publication.

### Identifying TE-compartmentalized genes

To identify TE-compartmentalized genes, for each species we first extracted the 50 kb flanking regions for each gene using bedtools (version 2.29.1) flank ^61^. We then intersected these regions with all coding regions (CDS) using bedtools intersect and filtered out flanking regions which overlapped CDS using bedtools subtract. We used bedtools coverage and species-specific TE annotations to calculate TE density for the noncoding flanking regions of each gene. We define TE density as the proportion of nucleotide positions occupied by TEs for a given region. We define TE-compartmentalized genes as genes within the top 90th (or 95th, see below) percentile of TE density when the full distribution of gene TE densities is considered independently for each species. We note that we explored multiple repeat density thresholds for TE compartmentalization, as well as different flanking region lengths (e.g. 10kb) and achieved similar results (Supplementary Tables 17-19). We also computed TE-compartmentalized genes separately for all TEs, LINEs, SINEs, LTR retrotransposons, and DNA transposons.

### Gene ontology database construction, analysis and filtering

We constructed species specific gene ontology (GO) databases for each genome. To do this, we extracted all gene ontology annotations corresponding to annotated genes in each genome from Uniprot ^57^ as well as AmiGo ^62^. Then, for each species, we filtered our gene set to only include genes with at least one GO annotation. We performed GO enrichment tests for TE-compartmentalized genes using goatools ^63^ with the flag *--method fdr_bh*. Goatools performs GO enrichment tests using Fisher’s exact tests with a false discovery rate correction. We required a minimum of four genes in test sets and filtered out species for which less than 20% of genes possessed GO annotations, resulting in the removal of 300 species (Supplementary Table 2). We first performed GO enrichment tests using TE-compartmentalized genes as defined by a 90th percentile repeat density threshold. In cases where we did not identify any enriched terms, we instead used a 95th percentile threshold. We performed these tests separately for all TEs, LINEs, SINEs, LTR retrotransposons, and DNA transposons.

Furthermore, after performing GO enrichment and purification (underrepresentation) tests for TE-compartmentalized genes in each species, we found that species which displayed no significant enrichment or purification results had significantly smaller GO databases than species with significant results (two-tailed ManwhitneyU test, P<0.0001). In light of this, we reasoned that our inability to find patterns of functional enrichment was likely due to poor GO annotation quality and/or lack of power in these species. Consistent with this, for species where we observed no GO terms whose p-values exceeded the FDR corrected threshold, the top 5 most enriched terms with p-values <0.05 showed consistent congruence with trends of term enrichment for species where we achieve FDR corrected significance. Thus, we considered these results in our downstream analyses. However, we filtered another 2 species which displayed no functional enrichment results with P<0.05. Together, after these filters, we were left with 732 species (Extended Data Table 2).

### GO Redundancy Filtration

To make sense of this complex dataset, we performed multidimensional scaling using a matrix of GO term semantic similarities through REVIGO, merging terms with dispensability > 0.5 ^64^. This reduced redundancy in our dataset and also provided metrics and a systematic framework with which to identify the most enriched terms and abundant terms. In our case, the dispensability metric is calculated based on both semantic relatedness to other terms and enrichment frequency. Figure 1B-D was produced using REVIGO output, with “Biological Process,” “Molecular Function” and “Cellular Component” terms labeled using dispensability cutoff of less than 0.0008, 0.04 and 0.02 respectively.

### Testing for molecular signatures of selection across gene families

We downloaded corresponding transcriptome fasta files from Refseq or Genbank for each considered species, and extracted the longest transcript for each coding gene in each genome. Then, we used a variation of the *FUSTr* pipeline ^65^ to cluster gene families and test for molecular signatures of positive selection. The *FUSTr* pipeline was developed for transcriptome data, and the primary purpose of the first four steps of the pipeline is transcriptome input processing. Thus, we skipped these steps. Briefly, the remaining steps of the pipeline are as follows. First, *FUSTr* performs an all-by-all blast between proteins using *diamond* ^66^ and clusters gene families using *silixx* ^67^. Then, *FUSTr* generates a multiple sequence alignment for each gene family using *MAFFT* ^68^ and a subsequent phylogeny using *fasttree* ^69^ as well as a trimmed alignment using *trimal* ^70^ for input to *FUBAR* ^48^, which performs tests for positive and negative selection at the molecular level. For downstream analyses and producing Figure 2, we required that each species have at least one TE-compartmentalized gene family and at least one non-TE-compartmentalized gene family.

### Genomic signature and chromosomal distribution analysis

Since GC-content can also contribute to differences in gene content as well as TE content within genomes, we also tested for differences in GC content between TE-compartmentalized genes and other genes across phyla. We separately calculated GC content for all genes as well as 50kb of noncoding sequence flanking genes using bedtools nuc ^61^. Although some individual eukaryotic lineages displayed differences in genic and/or gene-flanking GC content between TE-compartmentalized genes and other genes, we found very little evidence for any trend across diverse lineages (Extended Data Figure 3; two-tailed Binomial test, P=0.296). Recombination rates often correlate with TE distributions ^71^, and TEs can accumulate in regions of low recombination due to reduced efficacy of selection in such regions as well as selection against ectopic recombination in highly recombining regions ^72^. Thus, it is conceivable that reduced recombination rates for TE-compartmentalized genes could explain observed patterns. To evaluate this possibility, we intersected our TE compartmentalization results with recombination data for a subset of species (retrieved from ^73^). Specifically, we compared recombination rates for regions containing TE-compartmentalized genes to regions containing other genes. We find no significant trend of reduced recombination rates for TE-compartmentalized genes (Extended Data Figure 4; two-tailed Binomial test, P=0.151). Furthermore, for species in which TE-compartmentalized genes exhibit reduced recombination rates overall, we still observe outliers which display increased recombination rates relative to the median recombination rate for non-TE-compartmentalized genes (Extended Data Figure 4). Together, these results suggest that although recombination rates undoubtedly affects TE distributions, recombination rates alone fail to explain observed patterns

We explored chromosomal distributions of TE-compartmentalized genes for genomes with chromosome level assemblies. Specifically, we were interested in testing for enrichment or depletion of TE-compartmentalized genes at the ends of chromosomes (in prospective subtelomeric regions). Given the diversity of species considered in our study, most of which lack telomeric annotations, we crudely assigned subtelomeric regions based on chromosome size, using the first 10% and last 10% of bases on a given chromosome. Then, for each chromosome in each species, we compared observed proportions of TE-compartmentalized genes in these regions to expectations from random resampling across 1000 permutations.

### Population genetic analysis

We aligned 50 long-read based *D. melanogaster* genome assemblies to the *D. melanogaster* RefSeq assembly using minmap2 (version 2.21-r1071) with parameters -*a --eqx -x asm20 –cs -r2k -t 9* ^74^. Then, we used samtools (version 1.11) to convert alignments in sam format to sorted bam files ^75^. We employed svim-asm (version 1.0.3) with parameters *haploid --max_sv_size 100000000* to call genomic structural variants, SNPs and indels for each genome ^76^. We merged structural variant calls from each genome into one call set using SURVIVOR ^77^ with the parameters 500 1 1 1 0 50, which combines prospective redundant SV calls based on breakpoint vicinity (if both breakpoints are within 500 bp of each other) and filters for SVs longer than 50 bp. We extracted sequences for insertions and deletions, removed those which mostly contained ambiguous nucleotides, and ran Repeatmasker on them using our *D. melanogaster* TE library to identify TE-associated SVs. Finally, we filtered our structural variant calls for insertions or deletions which displayed at least 50% coverage for a TE. For SNPs and indels, we filtered for biallelic variants and merged calls for each genome using Bcftools (version 1.9) merge and Bcftools view respectively ^78^. We annotated synonymous and nonsynonymous variants with SnpEff (version 5.1d) ^79^. Population genetic analyses were performed using scikit-allel (version 1.3.3) (https://zenodo.org/badge/latestdoi/7890/cggh/scikit-allel).

### Comparing compartmentalization signatures between LTRs, LINEs and SINEs

We performed Fisher’s Exact Tests comparing proportions of TE-compartmentalized genes corresponding to each gene ontology term across all considered species between all combinations of LTR-compartmentalized genes, LINE-compartmentalized genes, and SINE-compartmentalized genes (Supplementary Tables 14-16). We used Q values to assess the proportion of terms with significantly different results for each combination of TE classes. This revealed that LINE-compartmentalized and LTR-compartmentalized genes showed significantly different enrichment results for only ∼31% of terms, whereas each significantly differed from SINE compartmentalized genes for >74% of terms. Since many GO terms are challenging to compare over long evolutionary distances, we also conducted these tests independently for different clades.

## Supporting information

Supplemental Tablel Captions

## Code availability

Code and documentation for our complete TE compartmentalization pipeline is available on github (https://github.com/lgozasht/TE-compartmentalization).

## Data availability

All data used in this work is publicly available on NCBI. All genomes considered for our TE compartmentalization pipeline are listed in Supplementary Table 1. *D. melanogaster* genomes used for population analyses are listed in Supplementary Table 13.

## Acknowledgements

The authors thank Cedric Feschotte, Sean Eddy, Tom Jones, Jenny Chen, Andreas Kautt, Olivia S. Harringmeyer, Matthew Hahn, James Mallet, Magnus Norborg and Timothy B. Sackton for helpful discussions and feedback on this manuscript. The authors thank Alexander Kramer for assistance with pipeline implementation. L.G. thanks Hopi E. Hoekstra for her support and advice throughout this work. The computations for this work were run on the FASRC Cannon cluster supported by the FAS Division of Science Research Computing Group at Harvard University. This work was supported in part by R35GM128932 to R.C.-D.

## Author contributions

L.G. and R.C.-D. conceived and designed the research. L.G. performed the research and analyses. All authors wrote the manuscript. All authors edited and contributed to the manuscript revision.

