## Extended Data Figures for "Universal signatures of transposable element compartmentalization across eukaryotic genomes"

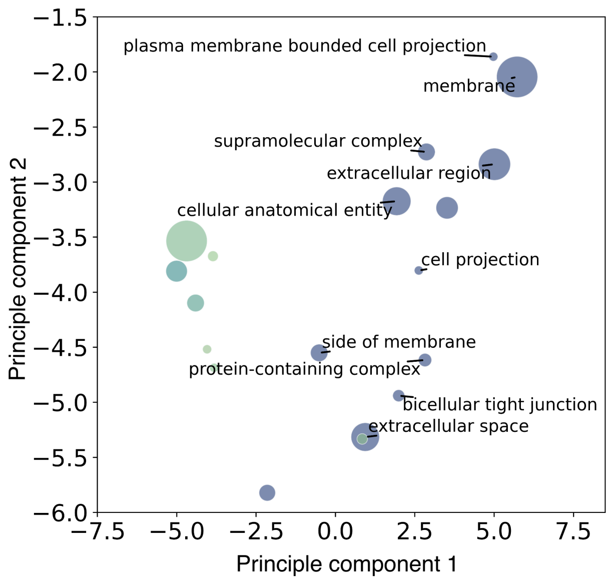


**Extended Data Figure 1:** Multidimensional scaling results for enriched Cellular Component gene ontology (GO) terms for TE-compartmentalized genes across all surveyed species (Supplementary Table 11). Each point represents a GO term, and point size and color correspond to enrichment frequency across species. Terms with dispensability metrics less than 0.02 are labeled to prevent overcrowding.

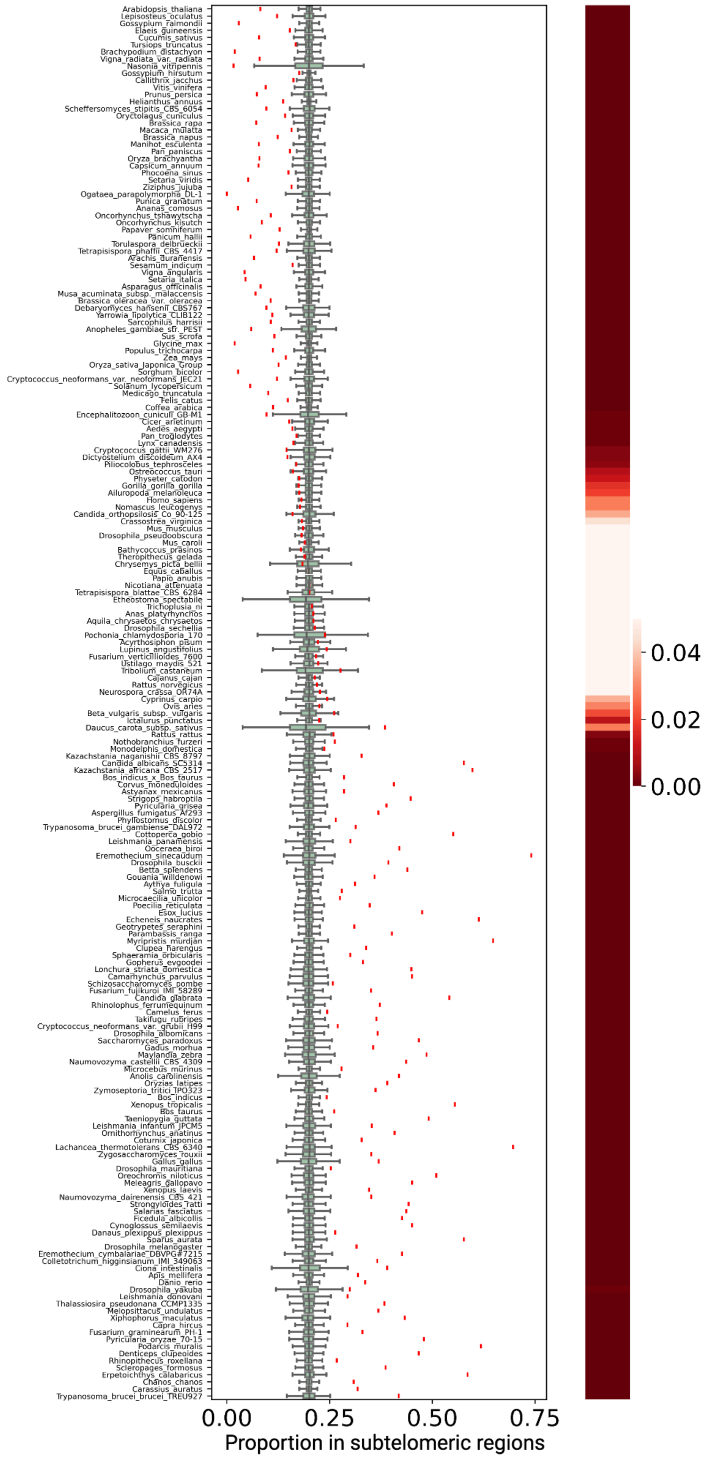


**Extended Data Figure 2:** Proportion of TE-compartmentalized genes in subtelomeric regions for each species with a chromosome-level genome as annotated by Refseq (see Methods). Red lines denote the observed proportion of TE-compartmentalized genes in subtelomeric regions and box plots represent expectations from randomization. Species are ordered by difference in mean expected and observed proportions, with species showing the strongest signatures of depletion for TE-compartmentalized genes in subtelomeres at the top. The heat map corresponds to P-values obtained from 1000 permutations comparing observed and expected proportions.

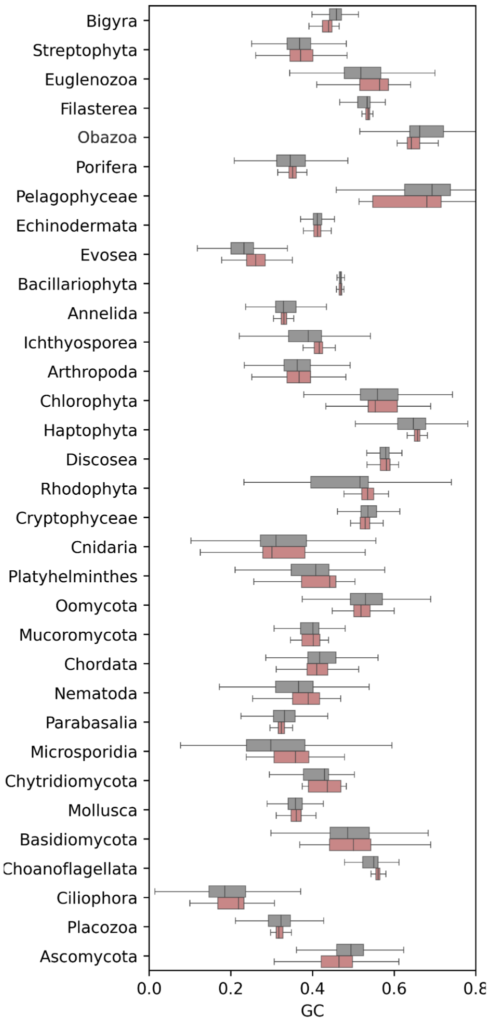


**Extended Data Figure 3:** Distributions of GC content for 5 kb regions adjacent to TE-compartmentalized genes (red) and other genes (gray) across diverse eukaryotic lineages.

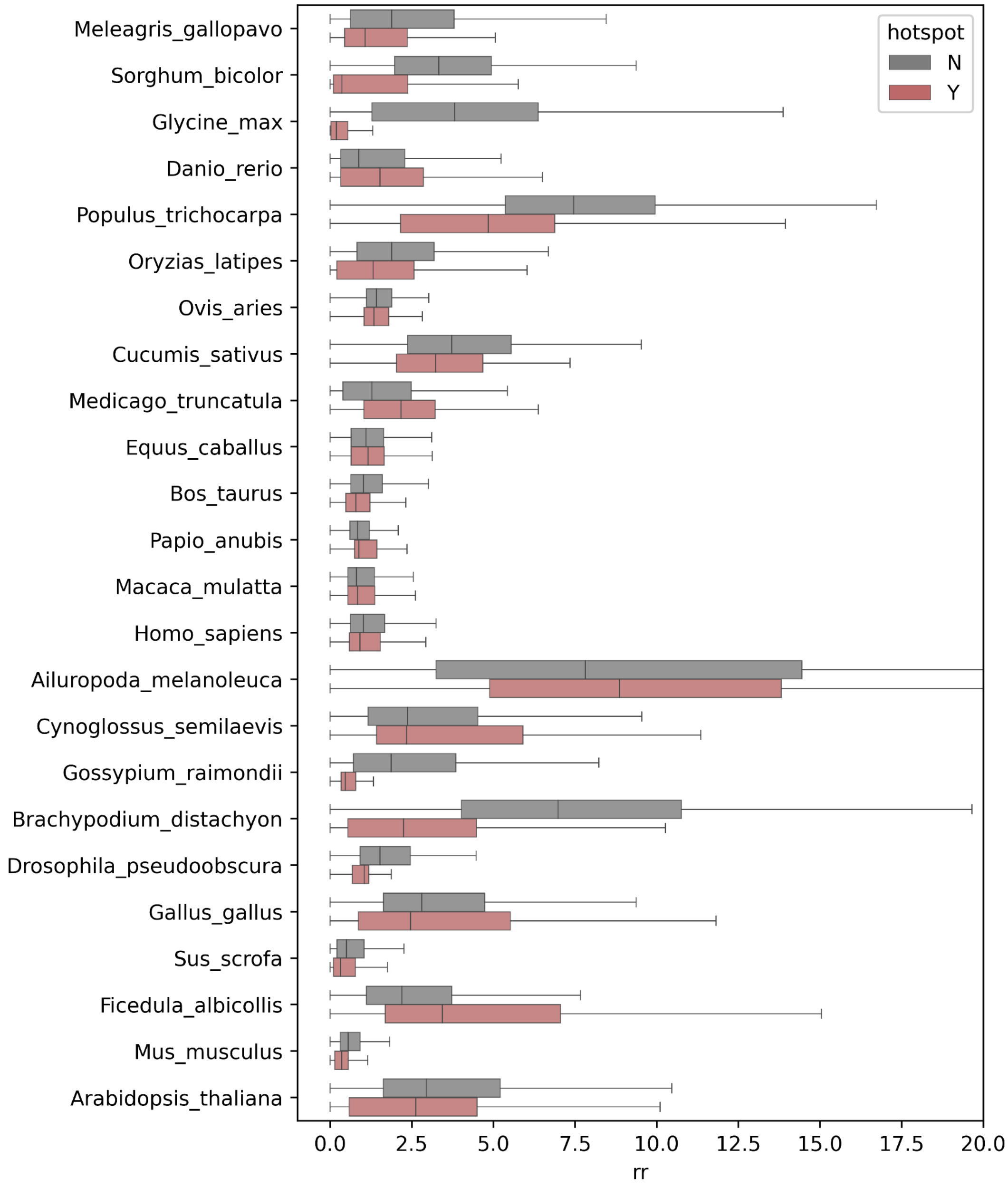


**Extended Data Figure 4:** Distributions of recombination rates for regions overlapping TE-compartmentalized genes (red) and other genes (gray).
