## Supplemental Tablel Captions for "Universal signatures of transposable element compartmentalization across eukaryotic genomes"

**Supplementary Table 1:** Refseq (GCF) and Genbank (GCA) genomes considered in this study.

**Supplementary Table 2:** Final set of genomes after filtering.

**Supplementary Table 3:** GO enrichment results for TE-compartmentalized genes in each species. Only terms with Q<0.05 are displayed for species in which at least one term shows Q<0.05. For species where no terms achieve Q<0.05 enrichment, the top 5 most enriched terms with P<0.05 are shown. For full enrichment results, see Supplementary Data 1.

**Supplementary Table 4:** GO enrichment results for TE-compartmentalized genes across specific TE classes in each species. Only terms with Q<0.05 are displayed for species in which at least one term shows Q<0.05. For full enrichment results, see Supplementary Data 1.

**Supplementary Table 5:** Summary of enrichment results for terms shared across species.

**Supplementary Table 6:** Summary of enrichment results for terms shared across species for specific TE classes.

**Supplementary Table 7:** Multigene family enrichment results for TE-compartmentalized genes and other genes.

**Supplementary Table 8:** Multigene family enrichment results for TE-compartmentalized genes and other genes across specific TE classes.

**Supplementary Table 9:** Multidimensional scaling of significant (Q<0.05) Biological Process GO enrichment results aggregated across species.

**Supplementary Table 10:** Multidimensional scaling of significant (Q<0.05) Molecular Function GO enrichment results aggregated across species.

**Supplementary Table 11:** Multidimensional scaling of significant (Q<0.05) Cellular Component GO enrichment results aggregated across species.

**Supplementary Table 12:** Proportions of sites showing signatures of positive and negative selection for each TE-compartmentalized gene family and each other gene family in each species.

**Supplementary Table 13:** NCBI accessions for long-read *D. melanogater* assemblies used for population genetic analysis.

**Supplementary Table 14:** Comparison of enrichment results for LTRs and SINEs across all GO terms. For each term (column 1), FDR corrected Fisher's Exact P-values (column 2) are calculated by comparing the proportion of LTR-compartmentalized to the proportion of SINE-compartmentalized genes corresponding to that term.

**Supplementary Table 15:** Comparison of enrichment results for LTRs and LINEs across all GO terms. For each term (column 1), FDR corrected Fisher's Exact P-values (column 2) are calculated by comparing the proportion of LTR-compartmentalized to the proportion of LINE-compartmentalized genes corresponding to that term.

**Supplementary Table 16:** Comparison of enrichment results for LINEs and SINEs across all GO terms. For each term (column 1), FDR corrected Fisher's Exact P-values (column 2) are calculated by comparing the proportion of LINE-compartmentalized to the proportion of SINE-compartmentalized genes corresponding to that term.

**Supplementary Table 17:** GO enrichment results for TE-compartmentalized genes using a 50 kb flanking window size and 95th percentile repeat density threshold. Only terms with $Q<0.05$ are displayed for species in which at least one term shows $Q<0.05$. For species where no terms achieve $Q<0.05$ enrichment, the top 5 most enriched terms with $P<0.05$ are shown.

**Supplementary Table 18:** GO enrichment results for TE-compartmentalized genes using a 10 kb flanking window size and 90th percentile repeat density threshold. Only terms with $Q<0.05$ are displayed for species in which at least one term shows $Q<0.05$. For species where no terms achieve $Q<0.05$ enrichment, the top 5 most enriched terms with $P<0.05$ are shown.

**Supplementary Table 19:** GO enrichment results for TE-compartmentalized genes using a 10 kb flanking window size and 95th percentile repeat density threshold. Only terms with $Q<0.05$ are displayed for species in which at least one term shows $Q<0.05$. For species where no terms achieve $Q<0.05$ enrichment, the top 5 most enriched terms with $P<0.05$ are shown.

**Supplementary Data 1:** Raw GO enrichment results for TE-compartmentalized genes in each species, including genes driving enrichment for each GO term.

**Supplementary Data 2:** Clade-specific GO enrichment result summaries.
